# The E3 ubiquitin ligase MIB-1 is necessary to form the nuclear halo in *C. elegans* sperm

**DOI:** 10.1101/301291

**Authors:** Leslie A. Herrera, Daniel A. Starr

**Author notes:** Correspondence to DAS: 1 Shields Ave Department of Molecular and Cellular Biology Davis, CA 95616.

## Abstract

Unlike the classical nuclear envelope with two membranes found in other eukaryotic cells, most nematode sperm nuclei are not encapsulated by membranes. Instead, they are surrounded by a nuclear halo of unknown composition. How the halo is formed and regulated is unknown. We used forward genetics to identify molecular lesions behind three classical *fer* (fertilization defective) mutations that disrupt the ultrastructure of the *C. elegans* sperm nuclear halo. We found *fer-2* and *fer-4* alleles to be nonsense mutations in *mib-1*. *fer-3* was caused by a nonsense mutation in *eri-3*. GFP::MIB-1 was expressed in the germline during early spermatogenesis, but not in mature sperm. *mib-1* encodes a conserved E3 ubiquitin ligase homologous to vertebrate Mib1 and Mib2, which function in Notch signaling. However, *C. elegans mib-1* has no Notch-related phenotypes. Thus, *mib-1* has been co-opted to regulate the formation of the nuclear halo during nematode spermatogenesis.

## 1. Introduction

The defining hallmark of eukaryotic cells is the nuclear envelope, a specialized extension of the endoplasmic reticulum consisting of two lipid bilayers and a perinuclear space (Hetzer, 2010). The essential role of the nuclear envelope is to compartmentalize the nucleus from the cytoplasm. Therefore, the nuclear envelope evolved in the last eukaryotic common ancestor and was critical to the evolution of the wide variety of eukaryotic organisms alive today (D. A. Baum and B. Baum, 2014). However, there are exceptions to this paradigm. All nematode classes except Enoplida have sperm devoid of a nuclear envelope at maturity (Justine, 2002; Yushin and Malakhov, 2014). Instead of two lipid bilayers, nematode sperm nuclei are surrounded by a halo of electron-dense material that also encapsulates the sperm centrioles (Wolf et al., 1978). The halo is thought to contain RNA (Ward et al., 1981). Nearly forty years after its initial discovery, little is known about the molecular makeup and developmental regulation of the perinuclear halo. We used a forward genetic approach in *C. elegans* to identify players in the formation of the perinuclear halo of nematode sperm.

*C. elegans* has long been appreciated as an excellent model to study gametogenesis (Hirsh et al., 1976). A large number of mutations have been isolated in *C. elegans* genes required for spermatogenesis, called *fer* or *spe* for <u>fer</u>tilization or <u>spe</u>rmatogenesis defective, respectively (Argon and Ward, 1980; L’Hernault et al., 1988). These mutations are specific to spermatogenesis and result in hermaphrodites that lay unfertilized oocytes. Many of these mutations are temperature sensitive. At the permissive temperature of 16°C, hermaphrodites lay normal, viable embryos, but at the restrictive temperature of 25°C, they lay almost 100% unfertilized oocytes. Moreover, the temperature sensitive period corresponds to the timing of spermatogenesis in hermaphrodites and the fertilization defect can be rescued by mating mutant hermaphrodites to wild-type males (Argon and Ward, 1980).

We were particularly interested in three genes, *fer-2*, *fer-3*, and *fer-4*, because of their striking ultrastructural mutant phenotype (Ward et al., 1981). When sperm from *fer-2*, *fer-3*, or *fer-4* mutant males grown at 25°C are examined by electron microscopy, the RNA halo that normally surrounds centrioles and condensed chromatin in mature spermatids is absent (Ward et al., 1981). In place of the halo, large tubules of straight hollow cylinders accumulate around the condensed chromatin of spermatids and sperm (Ward et al., 1981). The nature of the perinuclear tubules is unknown; with diameters of about 50 nm (Ward et al., 1981), they are unlike other described tubular cellular components. The working model, as proposed by Ward et al. (1981), is that in *fer-2*, *fer-3*, or *fer-4* mutant sperm, tubules form from aberrant polymerized components of the ribonucleoprotein complexes that normally make the perinuclear halo. We hypothesized that determining the molecular identity of *fer-2*, *fer-3*, and *fer-4* gene products would elucidate molecular mechanisms of the formation of the normal perinuclear halo and/or the abnormal tubules that form in the mutant sperm. Here we report that *fer-2* and/or *fer-4* is a mutation in the predicted E3 ubiquitin ligase *mib-1* and that *fer-3* is a mutation in *eri-3*, a member of a Dicer-associated complex.

## 2. Results and Discussion

We set out to identify the molecular lesions of *fer-2*, *fer-3*, and *fer-4* using a whole-genome sequencing approach. Libraries were prepared from *fer-2(hc2)* or *fer-3(hc3)* genomic DNA for Illumina paired-end sequencing. The sequences were aligned and analyzed using the CloudMap pipeline (Minevich et al., 2012) which identified over 5,000 single nucleotide polymorphisms (SNP) in the sequenced *fer-2(hc2)* strain when compared to the reference N2, wild-type genome. Only a single SNP in the *fer-2(hc2)* sequence data set was predicted to cause a nonsense mutation in an open reading frame. This SNP was a C to T transition at position 11,297,830 of chromosome III. It is predicted to change the tryptophan of codon number 460 in the *mib-1* gene to a stop codon. Thus, we hypothesized that *fer-2(hc2)* is an allele of *mib-1*.

We have high confidence in the *mib-1* mutation for the following three reasons. 1. The mutation was covered 105 times in the whole-genome dataset and confirmed by Sanger sequencing (Figure 1A). 2. Although *fer-2* was originally mapped to chromosome IV (Argon and Ward, 1980), three-point mapping later placed *fer-2* between *tra-1* and *dpy-18*, close to position 7.2 cM on chromosome III (Hodgkin, 1993). This position is within a cM of *mib-1*. 3. Based on the *fer-2(hc2)* phenotype, we predicted that *fer-2* transcripts would be enriched in the male germline. We therefore examined a list of 864 spermatogenesis-enriched genes previously identified in a microarray study (Reinke et al., 2004). *mib-1* is the only one of the 864 transcripts that maps between *tra-1* and *dpy-18* on chromosome III. Thus, *fer-2* maps near *mib-1* and *mib-1* is the only gene in the region enriched in spermatogenesis expression lists.

**Figure 1.**
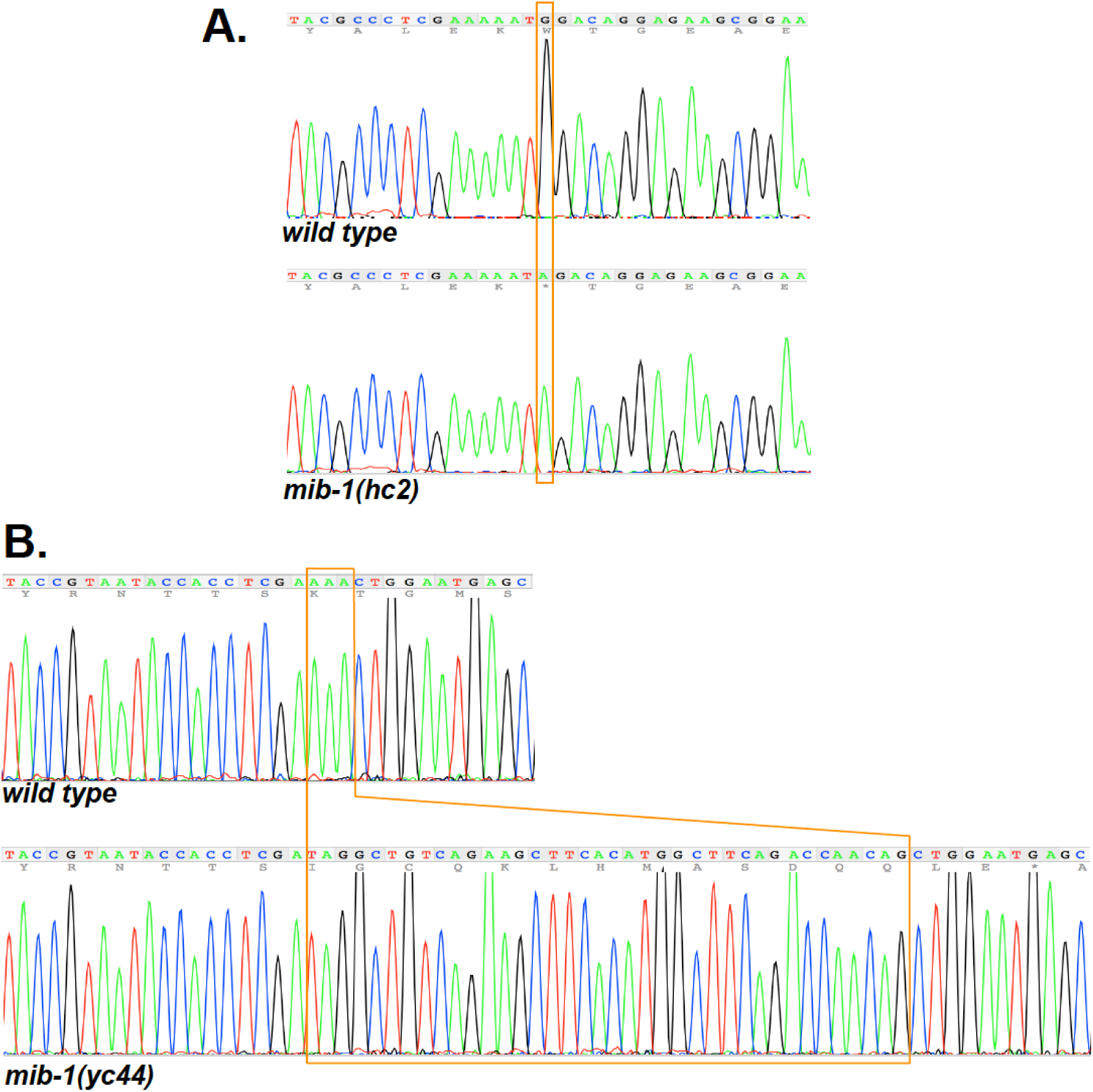
Mutations in *mib-1*. Sanger sequence outputs for wild type and *mib-1* mutant strains. (A) *hc2* is a C to T transition at position 11,297,830 of chromosome III. The opposite coding strand sequence is shown that leads to a Tryptophan to stop codon nonsense mutation. (B) *yc44* is a 38 bp insertion (boxed) in the sixth exon of *mib-1* that is quickly followed by a stop codon.

To further test whether *fer-2* is *mib-1*, one would traditionally attempt to rescue *fer-2(hc2)* with wild-type genomic DNA of the *mib-1* locus or to phenocopy the *fer-2(hc2)* phenotype with *mib-1(RNAi)*. However, extrachromosomal arrays are suppressed in germ lines and RNAi is very inefficient in *C. elegans* sperm. In addition, no known alleles of *mib-1* had been previously identified or characterized. We therefore used CRISPR/Cas9 to generate a new allele of *mib-1*. We isolated a mutant allele, *mib-1(yc44)* with a 38 base pair insertion of the *dpy-10* repair template that was used in our co-CRISPR approach (Arribere et al., 2014). The *yc44* insertion into the sixth exon of *mib-1* is predicted to cause a frame shift and is therefore likely a null allele (Figure 1B). *mib-1(yc44)* animals laid about 18% unfertilized oocytes when raised at 15°C, but laid an average of 97% unfertilized oocytes at 25°C (Figure 2). Thus, *mib-1(yc44)* phenocopied *fer-2(hc2)* as a temperature-sensitive spermatogenesis defective mutant.

**Figure 2.**
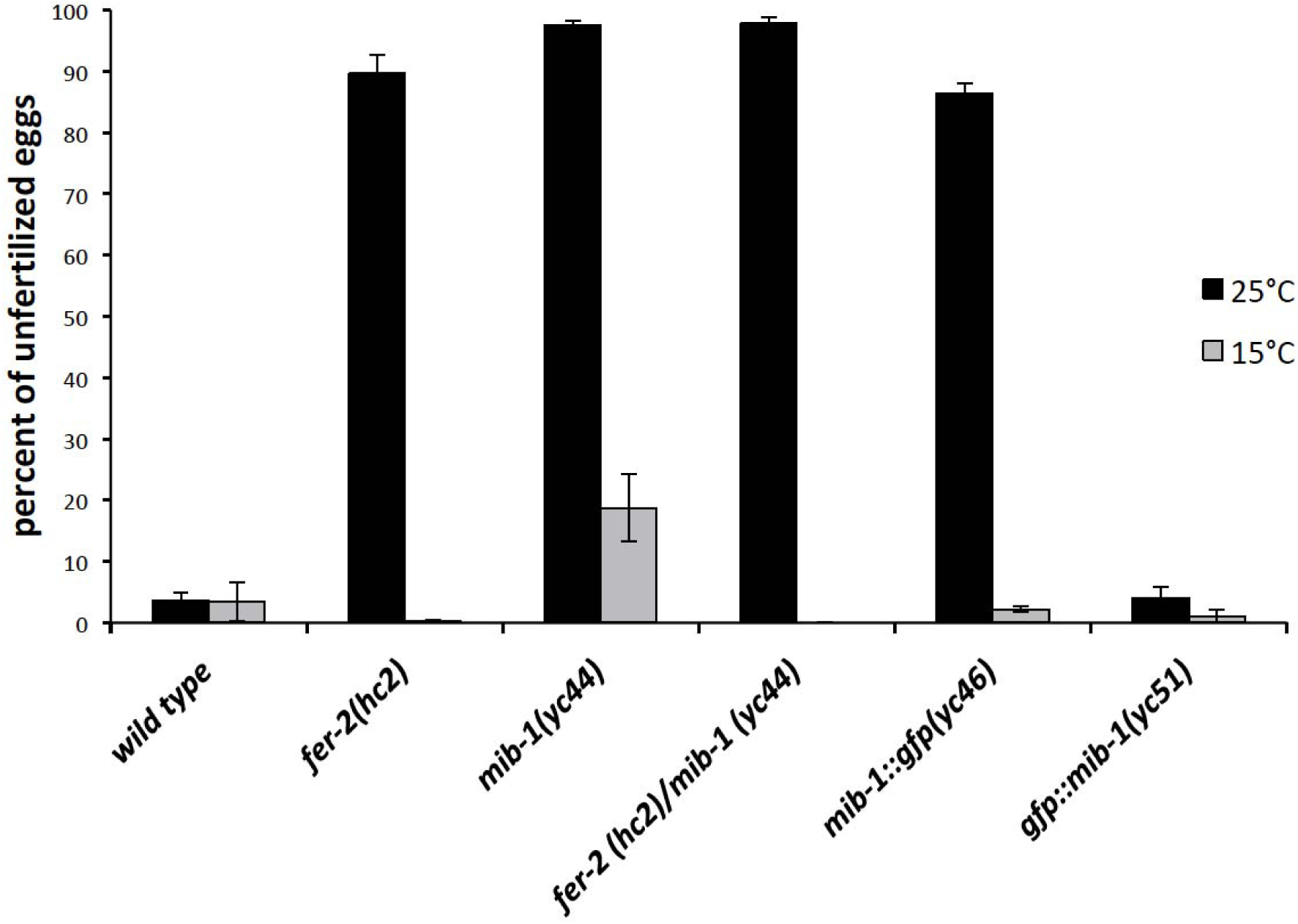
*mib-1* is required for male fertility. The mean percent of unfertilized oocytes from the total number of oocytes plus embryos laid by hermaphrodites is shown. For each bar, progeny from at least 10 hermaphrodites were scored at 15° and 25°C. Error bars are the standard error of the mean.

As final confirmation that *fer-2(hc2)* is an allele of *mib-1*, we crossed males homozygous for the C to T SNP on chromosome III to *mib-1(yc44)* hermaphrodites, and raised the F1 progeny at 25°C. The *fer-2(hc2)/mib-1(yc44)* heterozygotes laid 97.8% unfertilized oocytes at the restrictive temperature (Figure 2). Thus, *fer-2(hc2)* failed to complement *mib-1(yc44)* and we concluded that *hc2* is an allele of *mib-1*.

Our results suggest that one *fer-2(hc2)* strain has been mislabeled since its original isolation (Argon and Ward, 1980). We ordered all available strains carrying *hc2* or *hc4* from the Caenorhabditis Genetics Center and sequenced *mib-1* for the C to T mutation that causes the premature stop codon (Figure 1A) that we found in *hc2.* We identified the C to T mutation in BA2 *fer-2(hc2)*, BA4 *fer-4(hc4)*, and BA562 *fer-4(hc4)*; *him-5(e1490)*. However, the mutation was absent in BA547 *fer-2(hc2)*; *him-5(e1490).* Thus, an unidentified different lesion, perhaps outside of *mib-1*, likely causes the temperature-sensitive male sterile phenotype in the BA547 strain and it is not possible to determine whether we identified the original *fer-2(hc2)* or *fer-4(hc4)* lesion. We henceforth refer to both *fer-2* and *fer-4* as *mib-1*.

After determining the *fer-2* phenotype was due to a mutation in *mib-1*, CRISPR/Cas9 gene editing was used to tag the C-terminus of endogenous MIB-1 with GFP. Animals homozygous for *mib-1::gfp(yc46)* as the only source of MIB-1 lay an average of 86% unfertilized oocytes at 25°C (Figure 2), suggesting that MIB-1::GFP is nonfunctional. We proceeded to use CRISPR/Cas9 to tag MIB-1 with GFP at the N-terminus. *gfp::mib-1(yc51)* hermaphrodites laid wild-type percentages of fertilized embryos at both the permissive and restrictive temperatures (Figure 2), suggesting GFP::MIB-1 is functional. We next visualized GFP::MIB-1 by immunofluorescence with anti-GFP antibodies because of a weak GFP signal in live animals. GFP::MIB-1 was highly expressed in the proximal arm of the male gonad, but not in mature sperm (Figure 3). The timing of GFP::MIB-1 expression was consistent with a role in spermatogenesis, but not in mature spermatids; GFP::MIB-1 was turned off at about the time spermatids bud off of the residual bodies. GFP::MIB-1 was also seen in the L4 hermaphrodite gonad during spermatogenesis. GFP::MIB-1 was not detectable in other tissues.

**Figure 3.**
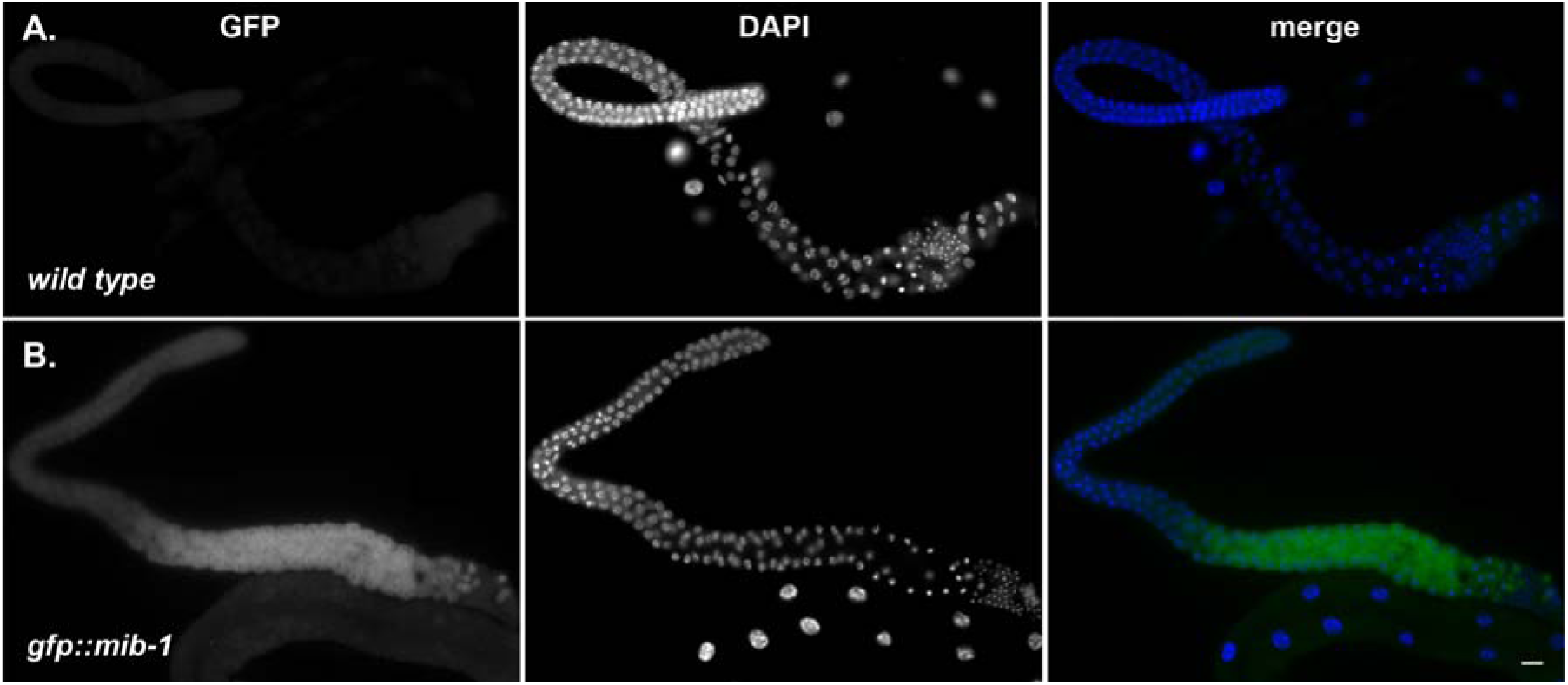
GFP::MIB-1 is expressed in the male germline. (A) wild type and (B) *gfp::mib-1.* Anti-GFP immunofluorescence showing GFP::MIB-1 expression in the proximal arm of male germline. GFP is green and DAPI-stained nuclei are blue in the merge. Scale bar is 10 μm.

*C. elegans mib-1* encodes an E3 ubiquitin protein ligase conserved to vertebrate Mib1 and Mib2 (Berndt et al., 2011). Mutations in the human *MIB1* gene cause left ventricular noncompaction cardiomyopathy (Luxán et al., 2013). Mib1 ubiquitinates the Notch ligands Delta and Jagged and targets them for endocytosis, turning off Notch signaling in zebrafish and mammals (Itoh et al., 2003; Kang et al., 2013). However, there are no major Notch-related phenotypes reported in *MIB2*^-/-^ knockout mice, suggesting that Mib2 acts synthetically to regulate Notch signaling and/or targets at least one alternate substrate (Guo et al., 2016; Koo et al., 2005). Our results show that *C. elegans mib-1* mutants have a fertilization defective phenotype that is distinct from any previously described Notch pathway mutant in *C. elegans* (Greenwald, 2005). Thus, it appears that *mib-1* has been co-opted in nematodes to function in perinuclear halo formation during spermatogenesis. The exact target(s) of MIB-1 in spermatogenesis remains to be determined.

We also cloned *fer-3(hc3)* by whole-genome sequencing. We had very little mapping data to guide us for *fer-3*. However, scanning the CloudMap output of SNPs between *fer-3(hc3)* and wild type led to the identification of a nonsense mutation in *eri-3*. We found a T to C transition at position 1,123,966 of chromosome II, which changes codon 69 of the *eri-3* gene from a serine to a proline. *eri-3* was previously reported to have a temperature sensitive spermatogenesis mutant phenotype (Pavelec et al., 2009). We crossed *fer-3(hc3)* into *eri-3(tm1361)* and the alleles failed to complement; F1 heterozygote hermaphrodites laid unfertilized embryos at 25°C. An additional group also reported that *fer-3(hc3)* is an allele of *eri-3* using the same complementation test (Conine et al., 2013). ERI-3 is a component of the 26G-RNA ALG-3/4 pathway and mutations in *alg-3/4* lead to a similar nuclear halo defect in spermatozoa, suggesting that they are in the same pathway (Conine et al., 2013). The expression of MIB-1 during spermatogenesis and a common downstream phenotype suggests a potential role in post-transcriptional regulation of proteins required for normal sperm morphology. It could therefore be informative in the future to identify the targets of MIB-1 and any possible relationship between MIB-1 and the 26G-RNA ALG-3/4 pathway.

## 3. Methods

Strains. *C. elegans* strains were grown on nematode growth medium plates spotted with OP50 bacteria and maintained at 15°C unless otherwise noted (Stiernagle, 2006). N2 was used as the wild-type control strain (Brenner, 1974). Strains were provided by the Caenorhabditis Genetics Center, which is funded by the National Institutes of Health Office of Research Infrastructure Programs (P40 OD010440). Strains BA2 *fer-2(hc2)*, BA3 *fer-3(hc3)*, BA4 *fer-4(hc4)*, BA547 *fer-2(hc2)*; *him-5(e1490)*, and BA562 *fer-4(hc4)*; *him-5(e1490)* were previously described (Argon and Ward, 1980). WM172 *eri-3(tm1361)* II was originally made by Shohei Mitani (National Bioresource Project at the Tokyo Women’s Medical University).

### Whole genome sequencing

For genomic DNA preps, nearly starved animals from five plates (5 cm each) were washed in M9 (Stiernagle, 2006), pelleted, resuspended in 200 μL buffer ATL (QIAGEN), and subjected to three freeze-thaw cycles. Genomic DNA was purified following the “animal tissue” protocol of the QIAGEN DNeasy Blood & Tissue Kit with an added RNAse A (10 mg/ml) incubation at room temperature for 15 minutes after the Proteinase K incubation. Genomic DNA was sent to the Functional Genomics Laboratory at QB3-Berkeley, fragmented into 300-600 bp pieces, and cloned into multiplexed libraries. The libraries were sequenced using paired end reads of 150 base pairs on an Illumina HiSeq 2500. Sequences were aligned, filtered and tabulated following the CloudMap pipeline (Minevich et al., 2012) on the public Galaxy server (Afgan et al., 2016).

### CRISPR/Cas9 mutagenesis and genome editing

CRISPR/Cas9 was used to generate a null mutation in *mib-1* and to fuse *gfp* to the 5’ or 3’ end of the *mib-1* open reading frame (Paix et al., 2015; 2016). Custom guide crRNA and universal tracrRNA were synthesized by Integrated DNA Technologies or Dharmacon. For the *mib-1(yc44)* deletion, the crRNA guide sequence was CGUAAUACCACCUCGAAAAC. For the *mib-1::gfp(yc46)* fusion, the sequence of the crRNA was AUUGAUAUUCACGAGUAGAU.

For *gfp::mib-1(yc51)* we used ACAAAAAUGAACGGAGUAGC as the crRNA. Purified Cas9-NLS protein was obtained from QB3-Berkeley. To make the repair templates for the homology-directed insertion, *gfp* sequences were amplified from pDD282 (a gift from Bob Goldstein; Addgene plasmid # 66823) (Dickinson et al., 2015) using Phusion DNA Polymerase (Thermo Fisher Scientific) and primers with overhangs consisting of 58-60 base pairs of *C. elegans* homology flanking the predicted CRIPSR/Cas9 cut-site (Paix et al., 2015). The QIAquick PCR Purification Kit (Qiagen) was used to clean the PCR product. In brief, we injected young adult hermaphrodite gonads with 9 μM or 17.5 μM crRNA:tracrRNA:Cas9 complexes along with 0.67 μM of the repair template for the *gfp* insertions (Paix et al., 2016; 2015). To follow CRISPR efficiency we used the *dpy-10* co-CRISPR approach (Arribere et al., 2014). Both a *dpy-10* crRNA sequence (1:12 ratio for *dpy-10* crRNA:*mib-1* crRNA) and 0.5 μM ssDNA oligo as a *dpy-10* repair template were added to the injection mix (Arribere et al., 2014). The *dpy-10* mutation was outcrossed to N2 or *him-8(e1489)* males after identification of the sought after genotypes by PCR screening. The following three strains were generated: UD549 *mib-1(yc44)*, UD563 *mib-1::gfp(yc46)*; *him-8(e1489)*, and UD577 *gfp::mib-1(yc51)*, *him-8(e1489)*.

### Fertilization and brood size phenotypes

To quantify the temperature-sensitive fertilization defects, assayed strains were initially cultured at 15°C on NGM plates seeded with OP50. Twenty L2 hermaphrodites from each strain were singled onto their own plates. For the *hc2/yc44* complementation test, we singled 40 F1 L2s expecting half would be males. Half of singled L2s were then raised at 25°C and the other half at 15°C. After 48 hours, hermaphrodites raised at 25°C were transferred to a fresh plate. The oocytes remaining on the original plate were scored as “fertilized” or “not fertilized” based on the globular and opaque appearance of unfertilized oocytes and the development of fertilized oocytes into L1 progeny within 24 hours. Oocytes laid within a 2-day window were added together. Plates with less than a total of 25 oocytes were excluded from the data. Also excluded were counts from F1s that were not cross progeny, which was verified by Sanger sequencing. For L2s raised at 15°C, counts started 72 hours after individuals were singled to account for the delay in development.

### Microscopy

We performed immunofluorescence staining on dissected male germlines from L4 males grown at 20°C. Dissection, fixation and immunofluorescence were performed as described (Jaramillo-Lambert et al., 2007) with the following alterations. Samples were fixed for 5 minutes at room temperature in 2% paraformaldehyde in egg buffer [118 mM NaCl, 48 mM KCl2, 2 mM CaCl2, 2 mM MgCl2, 5 mM HEPES at pH 7.4]. We used a 0.7% BSA/1× PBS + 0.1% Tween20 solution for blocking. Rabbit anti-GFP antibody (NovusBiologicals NB600-308) was used at a 1:500 dilution and donkey anti-rabbit antibody Alexa Fluor 488 (Invitrogen A21206) was used at 1:500 for the secondary. DNA was stained with DAPI (final concentration of 0.2 ng/uL).

Images were collected with a 63× Plan Apo 1.40 NA objective on an DM6000 epifluorescence compound microscope (Leica) with AF6000 software (Leica). Images were uniformly enhanced using the levels command in Adobe Photoshop.

## Acknowledgements

We thank David Fay and members of the Fay lab for hosting DAS on sabbatical where the project was initiated, especially John Yochem who helped analyze the whole-genome sequences. We thank QB3 Berkeley for preparing libraries and Illumina sequencing. We thank Venecia A. Valdez and Michael Paddy for help with the imaging. We thank members of the Starr lab for reading the manuscript and support throughout the project. This work was supported by the National Institutes of Health (grant number R01 GM073874).

